# Biomimetic system design for engineering biofidelic 3-D respiratory tissues *in vitro*

**DOI:** 10.1101/2020.06.25.172700

**Authors:** C. Poon, M. Zhang, P. Boughton, A. Hong, A. Ruys

## Abstract

**Objective:** The structural and functional complexity of the respiratory system present significant challenges to capturing conditions vital for maintaining phenotypic cellular functions in vitro. Here we report a unique tissue engineering system that enables respiratory constructs to be cultured under physiological loading at an air-liquid interface (ALI).

**Methods:** The system consists of a porous poly-e-caprolactone scaffold mounted in a well insert, which articulates via magnetic coupling with a linear actuator device to strain attached scaffolds through a sterile barrier. For proof of concept, NCI-H460 human carcinoma cells were seeded on scaffold inserts which were subjected to 5-15% cyclic tensile strain at 0.2Hz within a six well plate. The dynamic constructs were cultured at an ALI in a standard incubator for up to 10 days along with unstimulated (static) ALI and static submerged control groups.

**Results:** High (near-100%) cell seeding efficiency was achieved within the scaffold-strain device. Both dynamic and static ALI groups yielded higher cell densities compared to the submerged control for all time points. Distinctly different patterns in cellular growth and behaviour between dynamic air-liquid interface and conventional static submerged culture groups were revealed by nuclei staining, where the actuated group displayed more uniform cellular distribution throughout the construct compared to both static controls.

**Conclusion:** Air-liquid interface culture and physiological strain are important for engineering respiratory tissue models.

**Significance:** The system described allows scalable and replicable culture of 3-D tissue engineered respiratory models under biologically-relevant conditions.

## I. Introduction

The conventional approach to *in vitro* cell culture uses two-dimensional (2-D) surfaces to attach and grow cells on tissue culture polystyrenes. While central to much of our current research paradigm, it is well established that 2-D culture environments fail to reproduce many of the cell-cell signaling and external cues experienced by cells *in vivo* which are vital to their function [1, 2]. Cell behaviors under 2-D conditions vary significantly from those in 3-D across a range of properties including cell morphology, gene profile and migration behavior [3-5]. Critically, de-differentiation and the restricted life spans of cells in 2-D culture limit their predictive potential and make it difficult to undertake long-term studies [6-8]. These issues, in addition to growing ethical and scientific concerns regarding the use of animal models [9, 10], have motivated a paradigm shift towards 3-D tissue culture approaches which aim to deliver critical conditions experienced by cells *in vivo* that conventional 2-D culture techniques intrinsically lack.

A popular approach to 3-D tissue culture is a scaffold-based construct, where synthetic and natural materials are used to create a supportive matrix for cell attachment and growth [11]. More recently, increased understanding of the critical role of mechanical stimuli in modulating cellular functions has prompted development of *in vitro* culture devices that aim to replicate tissue and organ-specific biomechanical conditions [12-14]. Such systems may employ perfusion or direct mechanical actuation to stimulate a 3-D construct [12, 15-19], where the method of delivering stimulus and system design are determined by model and application specific requirements [20, 21]. Several strategies for respiratory tissue engineering have been described, including decellularized ECM scaffolds [22-24], air-liquid interface well inserts [25-27] and PDMS microfluidic chips with integrated 2-D membranes [27-29]. However, biologically-derived materials introduce batch-batch inconsistencies and there exists a need for a truly 3-D *in vitro* culture model that also provides mechanical stimuli.

Our research investigates critical biological and engineering principles required for producing biofidelic, reproducible and scalable 3-D *in vitro* respiratory tissue models. In this study, we present a proof-of-concept follow up to our previous work which introduced an *in vitro* system designed to deliver key physiological and biomechanical respiratory tissue conditions to 3-D cultures on an optimized porous synthetic scaffold [30]. Here, we demonstrate that a 3-D lung tumor model cultured under ALI and physiological strain conditions within our unique *in vitro* scaffold-strain system display distinctly different cellular behaviors compared to static submerged controls, affirming the underlying biomimetic design rationale. Overall, our results affirm the necessity of recapitulating physiological respiratory conditions for promoting optimal cellular functions *in vitro* and represent an advancement towards engineering biologically-accurate respiratory tissue models.

## II. Capturing respiratory cellular conditions in vitro

The complex structural, functional and biomechanical characteristics of the human lung present significant challenges to simulating the cellular conditions experienced by the alveolar epithelium *in vitro*. Lung tissues are highly elastic and porous, consisting of an interconnected network of alveolar sacs that undergo periodic strain and relaxation. During each breath, these sacs are recruited along a multidimensional pressure gradient within a range of magnitudes that depend on their location within the lung, tissue condition and breath volume [31]. Accordingly, the macroscopic strain profile of lung tissue follows two distinct phases: inspiration and expiration.

During inspiration, active contraction of diaphragm causes the volume of the lung and total surface area of the respiratory epithelium to expand. As the epithelium is strained, alveolar cells flatten to minimize the thickness of the blood-air-barrier, thereby facilitating gas exchange. During expiration, passive relaxation of the diaphragm and elastic recoil of the alveolar ECM compress the alveoli, and cells are remoistened by extracellular fluids. This cyclical biphasic loading regime, in concert with periodic aeration and wetting of the respiratory epithelium, is characteristic of the respiratory environment and must be recreated to accurately model distal lung tissue conditions *in vitro*. Table 1 summarizes key *in vivo* properties and conditions found within the lung, which formulate design parameters required for more biologically-relevant culture of lung tissues.

**Table 1.** Biomimetic design parameters determined by physiological conditions and properties of the lung

| Parameter | In vivo values | System values | References |
| --- | --- | --- | --- |
| Tissue strain | 5-15% | 5-15% | [32, 33] |
| Breathing rate (at rest) | 12-15 breaths/minute | 12 cycles/minute (0.2 Hz) | [34] |
| Inspiration: expiration rate | 1:1.5-2 seconds | 1.6:3.3 (seconds) | [35] |
| Alveolus diameter | 200-300 $\mu\text{m}$ | Scaffold pores 200-300 $\mu\text{m}$ | [36] |
| Temperature | 37±0.5 $^{\circ}\text{C}$ | 37±0.5 $^{\circ}\text{C}$ | N/A |
| O<sub>2</sub> (inspired-expired) | 20%-15% | 20% | [34] |
| CO<sub>2</sub> (inspired-expired) | 0.03%-3.6% | 0.03%-5% |  |
| Oxygen tension | Blood-air barrier | Air-liquid interface | [37, 38] |

Our device achieves these conditions by delivering physiological strain to a structurally-relevant scaffold at an air-liquid interface (ALI). The system is comprised of a linear actuator designed to mechanically stimulate up to six replicates within a standard disposable six well culture plate. The actuation mechanism utilizes a programmable servo motor to drive a bed of six permanent magnets which couple with individual magnets embedded in strain-transducing well inserts through the base of the tissue culture plate. Scaffolds or samples are fused to the well inserts and cultured within a standard six well plate (Figure 1). Magnetic interaction between the inserts and linear motion of the magnet bed induces controlled actuation of the cultured constructs across a sterile barrier.

**Figure 1.**
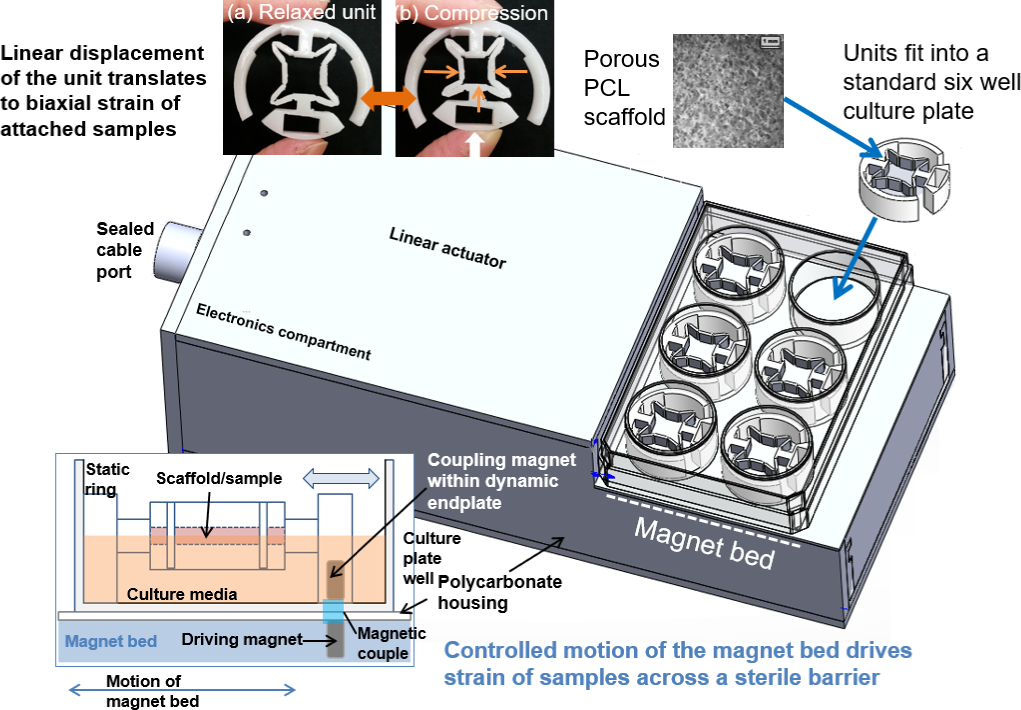
The linear actuator-driven system delivers mechanical stimuli to constructs cultured in a standard culture plate via magnetic coupling. Insets (a) & (b): an articulating well insert translates linear motion to axial and transverse strain across attached scaffold cultures at an air-liquid interface; bottom: side schematic view of the insert mechanism

The system operates in a standard incubator for controlled temperature, humidity and gas composition. A unique scaffold well insert was specifically developed to model strain conditions native to the lung. The insert mechanism consists of a hinged frame that translates a 3 mm linear displacement of the magnetic endplate to 5-15% strain gradient across the axial and transverse axes of a 11 x 11 x 3 mm porous polycaprolactone scaffold, on which cells are seeded and cultured (Figure 1). Compared to our previously introduced design [30], this open configuration achieves consistent axial and transverse strain, high cell seeding efficiency as well as facilitates extraction of the scaffold for analysis and potential live cell imaging.

## III. Methods & Materials

### A. Fabrication

97% (w/w) porous scaffolds were fabricated from polycaprolactone (CAPA 60000, Perstorp, Sweden) via a combination of solvent casting and particulate leaching methods using a sucrose porogen to template pore sizes of approximately 200-300μm, the diameter of a typical alveolus [18]. Inserts were fabricated using the same polycaprolactone via rapid prototyping and custom silicone moulds. Scaffolds were fused to the inserts with a custom polycaprolactone-based adhesive to avoid the cytotoxicity of cyanoacrylate-based adhesives. Scaffold inserts with heights of 11 mm and 7.5 mm were fabricated to compare air-liquid interface (dynamic and static) and submerged culture conditions respectively. Assembled scaffold inserts were sterilized by rinsing with 70% ethanol and stored dry in six well culture plates.

### B. Experimental Setup

Inserts were rinsed with 70% ethanol and phosphate buffer saline solution (PBS), UV irradiated for 20 minutes, dried in a fume hood for 24 hours then transferred to new six well culture plates. A near-confluent T75 flask of H460 human lung carcinoma cell line subcultured in 5% FCS supplemented RPMI-1640 medium (Gibco Life Sciences) was harvested and made up to a concentration of 5 × 10^6^ cells/mL. 100µL of seeding stock solution was pipetted at the center of each scaffold to seed 5 × 10^5^ cells/scaffold. After seeding, the inserts were incubated at 37 °C for 60 minutes to allow cell attachment. 6 mL of 10 % FCS supplemented RMPI medium were then added to each well such that the meniscus occurred at the upper plane of the ALI scaffolds (Figure 2). All seeded scaffold inserts were incubated statically for 24 hours allow cell attachment to the scaffolds, then transferred to new plates with fresh media. The cell seeding plates were stained with Coomassie Brilliant Blue dye solution to visualize the presence of detached cells as an indicator of seeding efficiency.

**Figure 2.**
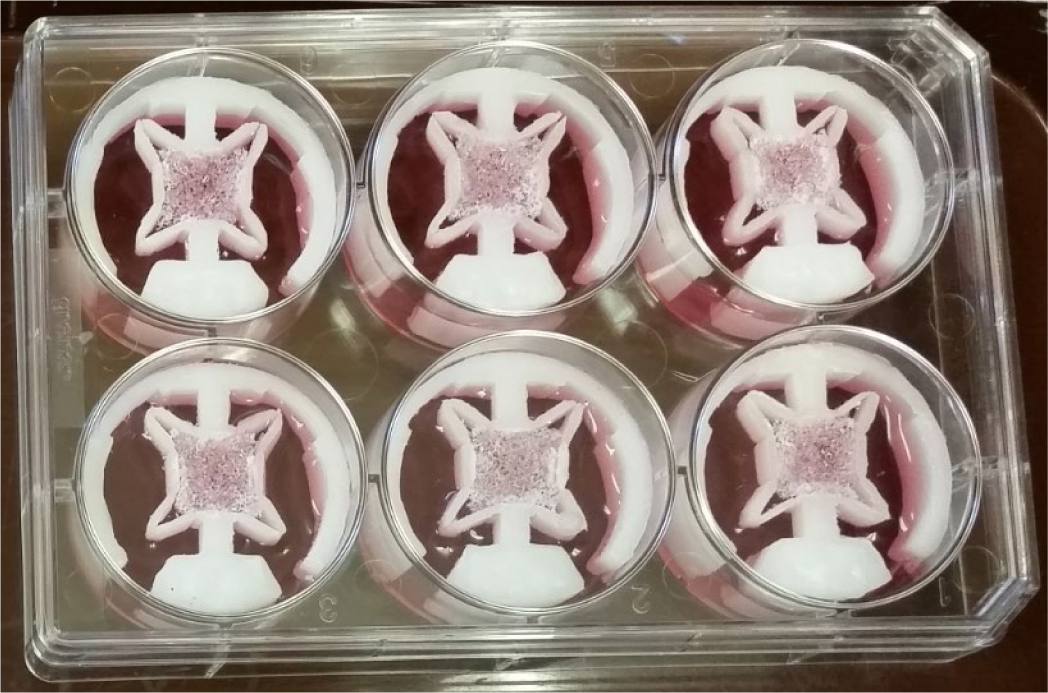
Seeded scaffold inserts in a standard six well culture plate

After 24 hours, the dynamic ALI group was placed on the magnetic actuator and strained at 0.2 Hz with 3 mm linear displacement (equating to 5-10% axial and transverse strain of the scaffold) within a standard incubator. Both control groups were incubated statically. A pair of scaffold inserts from each group were harvested in 72 hour increments at 4, 7 and 10 days of culture. Unharvested inserts at each time point were transferred to new six well plates with fresh culture medium. Total protein analysis and nuclei staining were performed to quantify cell proliferation across the 3-D constructs and visually assess and compare cellular distribution under the different culture conditions respectively.

### C. Nuclei Staining for Visual Analysis

Harvested scaffold inserts were rinsed with PBS in a six well plate then submerged in 1.5% methylene blue solution (Sigma Aldrich) for 30 minutes at room temperature, then de-stained by rinsing thrice with deionised water. Stained inserts were stored in a fresh six well plate and refrigerated until all time points were obtained. Given the presence of cell colonies on the strain mechanism and insert arms, entire scaffold inserts were imaged macroscopically on a cell colony counter (Bantex 920A) to capture a more comprehensive representation of cell growth, distribution for a more holistic assessment of cellular behaviors.

### D. Total Protein Analysis

Total protein concentrations from cell lysate obtained from the scaffolds at 4, 7 and 10 days of culture were assayed to provide a direct quantitative measure of cell number and thus inform proliferation trends within the 3-D constructs.

#### 1) Sample preparation

Harvested inserts were gently rinsed with PBS in a six well plate and scaffolds were extracted from the strain mechanism using a scalpel blade. To remove traces of culture media proteins, the scaffold and PBS from each sample were transferred to capped 10mL tubes and centrifuged at 750 rpm for 3 minutes. The supernatant was removed and the process was repeated with fresh PBS. The scaffold and cell pellet was frozen with 300µL cell lysis buffer (#9803, Cell Signaling Technology, USA), then thawed and refrozen thrice to ensure complete cell lysis. Samples were then transferred to 2mL Eppendorf tubes, kept on ice and sonicated thrice in 20 second intervals (QSonica). Sonicated samples were then centrifuged at 2000 rpm for 2 minutes, and the supernatant was kept frozen in 1 mL Eppendorf tubes until all time points were obtained for total protein analysis.

#### 2) Protein concentration assay

96 well microplates were preloaded with 200 µL 20% Coomassie Brilliant Blue dye solution (#500.0006, BioRad®) made up with deionized water. A standard curve was established with bovine serum albumin (BSA) concentrations of 0, 0.05, 0.1, 0.2, 0.3, 0.4, 0.5 and 0.6 mg/mL. Samples were thawed and thoroughly mixed in a tube shaker for 10 seconds, then diluted to 1:10 with deionized water in 0.5mL Eppendorf tubes with further dilution as necessary. The protein standards and samples were then loaded in prepared microplates and absorbances were read at 595 nm (EnSpire® Multimode Plate Reader™). Raw absorbance data was exported as for analysis. A standard curve was established from protein standard absorbance values, and total protein concentrations of the samples were then calculated using the standard curve and respective dilution factors. The assay was repeated in triplicate.

## IV. Results & Discussion

### A. System operation

The linear actuator operated reliably over continuous periods of 10 days, providing over 72,000 actuation cycles per assay. The magnetic coupling between the driving endplate and well insert was robust, providing controlled displacement. No wear debris or scaffold disintegration was observed during routine checks of the culture plate wells at each harvesting time point, indicating the suitability of the polycaprolactone scaffolds and inserts for prolonged dynamic culture.

### B. Cell seeding efficiency

The 100µL bolus of cell seeding solution was retained within each scaffold for all samples. There was no evidence of leakage as previously experienced with static submerged scaffold culture during scaffold development. Leakage results in low seeding efficiency and inconsistent seeding, which in turn leads to results variability to variable and unknown numbers of cells present within each construct *de novo*. Protein coatings on culture plates may also influence substrate preference and impact seeding efficiency at contacting surfaces.

Cell smears are clearly evident on the base of the wells of scaffolds that were dry-seeded then cultured in a submerged free-floating condition (Figure 3A) these are cells that have detached or migrated off the scaffold after seeding, leading to a low seeding efficiency estimated to be ∼10% during scaffold development and optimisation. In contrast, few isolated cells can be observed on the bottom of the scaffold insert wells 24 hours after seeding (Figure 3B). While not a conventional means of quantifying seeding efficiency, the lack of cells on the culture plate after seeding indicate that nearly 100% of the seeded cells were successfully retained within the scaffold. This was achieved by suspension of the porous scaffold within the strain mechanism of the insert and addition of a smaller volume of seeding medium than the volume of the scaffold. This bolus is effectively retained by surface tension within the scaffold and so the only surfaces available for cells to attach and establish on are limited to the scaffold, resulting in high cell seeding efficiency.

**Figure 3.**
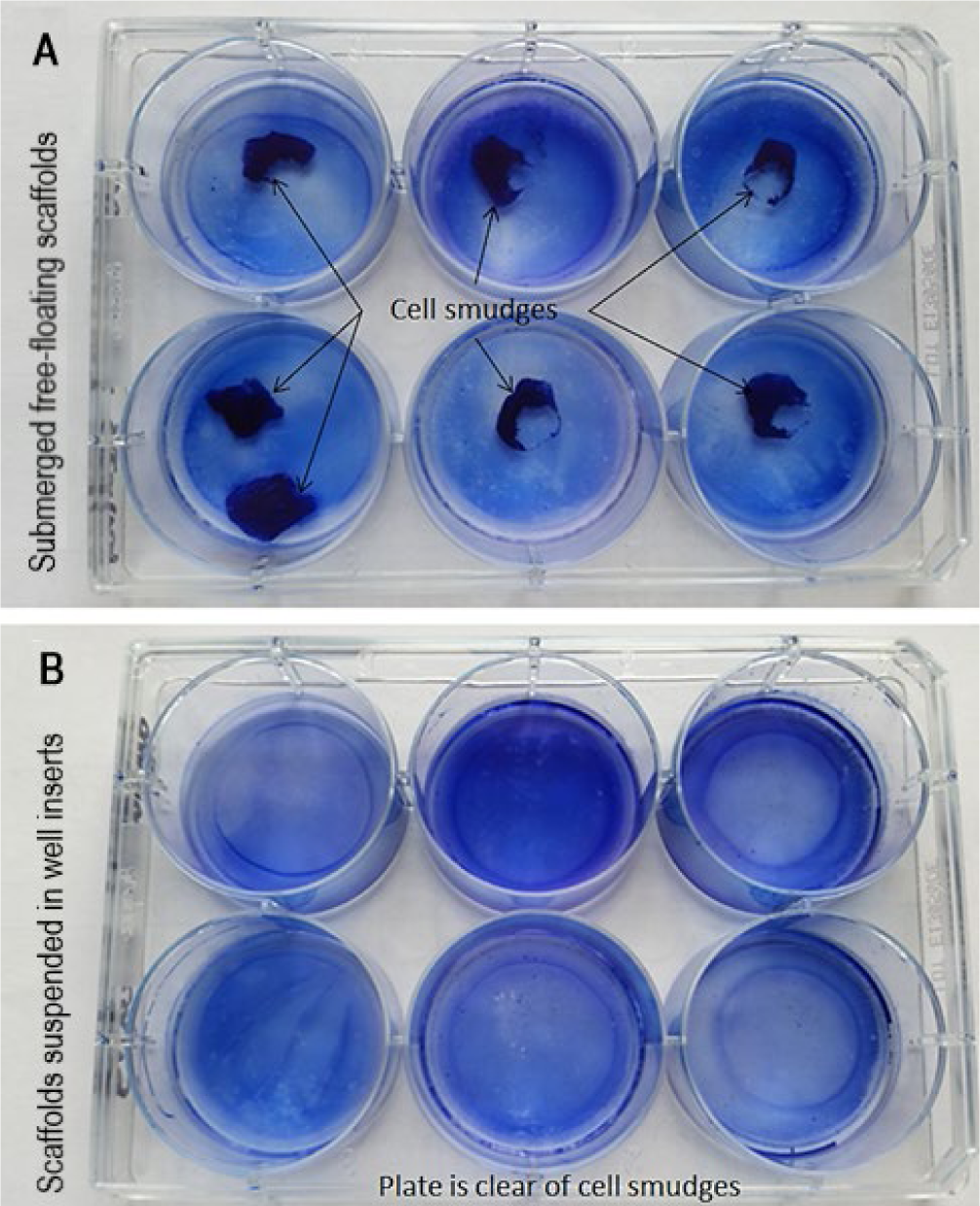
Culture plates stained with Coomassie Brilliant Blue dye after passive cell seeding and incubation for 24 hours. A. Cell smears from free-floating scaffolds; B. Cell-free wells after suspension-seeding scaffolds in well inserts

As the lack of cells present on the seeding plate is directly correlated to cell retention within the scaffold, this result represents an improvement to conventionally low seeding efficiencies of ∼10-25% reported for passive seeding of 3-D porous scaffolds [39]. Given that no specialized techniques such as centrifugation, vacuum suction, bioreactor-mediated seeding, or additional steps were necessary to achieve this level of cell retention, such a simple suspension culture approach is recommended or seeding porous 3-D scaffolds.

### C. Visualizing cell growth patterns in 3-D constructs

#### 1) Nuclei staining

Methylene blue (nuclei) staining was found to be effective for visualizing the presence and distribution of cells cultured on the scaffold inserts (Figure 4, Figure 5). The stain provided a direct correlation between staining intensity and cellular density i.e. more deeply stained regions on the scaffolds are indicative of higher cell numbers, and vice versa. The polycaprolactone scaffolds and insert components were discolored but not stained by methylene blue (Figure 4, Figure 5), providing an effective contrast to visualizing the cells. Conventional means of assessing cellular infiltration into scaffolds include cryosectioning and histological analysis using standard microscopy techniques. However, macroscopic imaging provides valuable information on cellular growth, proliferation and migration patterns across the whole system. Studying whole inserts rather than scaffolds (or sections) in isolation is important as different substrate surfaces (the scaffold, strain mechanism and culture plate) and conditions represent different environments which determine cellular behavior, intra-cellular signaling and construct remodeling. Therefore, variations in cellular growth and distribution patterns between experimental groups represent key indicators of different cellular behaviors induced by respective constituent culture conditions. Thus, nuclei staining and macroscopic imaging enables more holistic design evaluation and is recommended in conjunction with standard microscopy.

**Figure 4.**
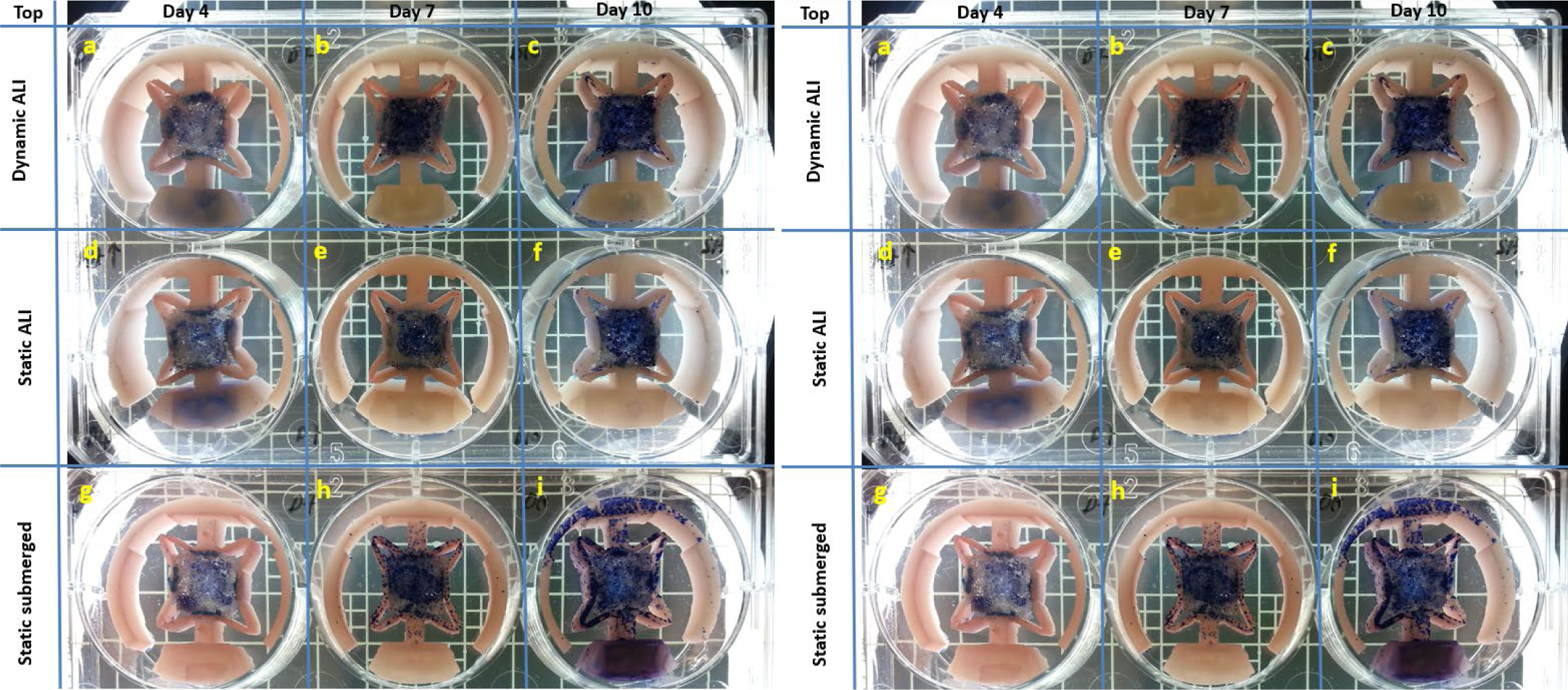
Top (left) and base (right) of the scaffold inserts from all experimental groups and time points after staining with methylene blue dye as viewed on a cell colony counter. Cells were stained blue, while the polycaprolactone of the scaffold and units remained unstained/turned light pink

**Figure 5.**
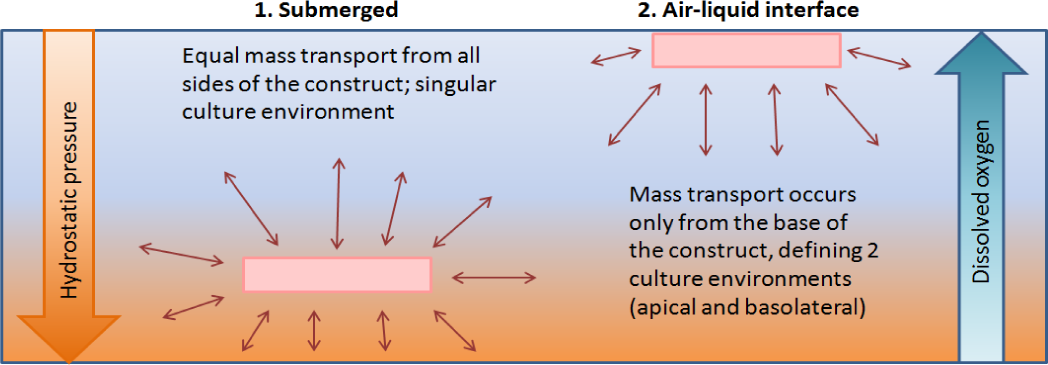
Comparison of submerged and air-liquid interface culture conditions at different depths within a culture well

#### 2) Cell growth & proliferation

NCI-H460 cells established on and progressively filled the polycaprolactone scaffolds across the 10 day culture period for all groups as reflected in the chronological changes in staining intensity shown in Figure 4 and Figure 5. Proliferation of cells throughout the scaffold confirm high pore interconnectivity, sufficient mass transport and adequate physical cues in promoting adhesion, migration and growth [40, 41]. Taking staining intensity as a direct indicator of cell numbers, the most complete and even cellular distribution across the scaffold was found to occur in the actuated ALI groups at day 7 and 10 (Figure 4, Figure 5). In contrast, cells grew as clustered colonies amid unstained regions of bare scaffold in both statically cultured control groups, indicating lower cellular infiltration. As cellular migration and infiltration within a 3-D construct are key indicators of phenotypic function [13, 42], particularly relevant to metastatic cell lines such as NCI-H460, this result confirms that mechanical stimuli provided by the scaffold strain device has a positive effect on cellular behavior. Staining intensity (cell density) markedly increased between days 4 and 7 for all groups. At day 7, the actuated ALI group was the most densely stained (contained the highest cell density) whereas the static submerged scaffolds were the least stained, with significant numbers of off-scaffold cells present on the strain mechanism (Figure 5). Cell numbers appeared to remain similar between days 7 and 10 for both ALI groups or slightly decreased for the static submerged control group. This can potentially be attributed to: (i) the cell population reaching saturation within the scaffold, (ii) the formation of a necrotic core due to outer cell layers impeding mass transport through the scaffold, or (iii) general plateauing of proliferation after reaching a stationary density [43, 44]. The effect of initial seeding density on construct remodeling warrants further investigation.

#### 3) Cellular response to culture environment

Each experimental group was found to display different patterns in growth and cellular distribution within the scaffold inserts (Figure 4, Figure 5), suggesting that within the range of conditions present in a 3-D culture system, cells preferentially respond to a unique subset of conditions within this range closest to those experienced *in vivo* as summarized in Figure 6. In addition to previously discussed construct growth trends, the location and density of off-scaffold cells indicate different cellular behaviors between the experimental groups.

**Figure 6.**
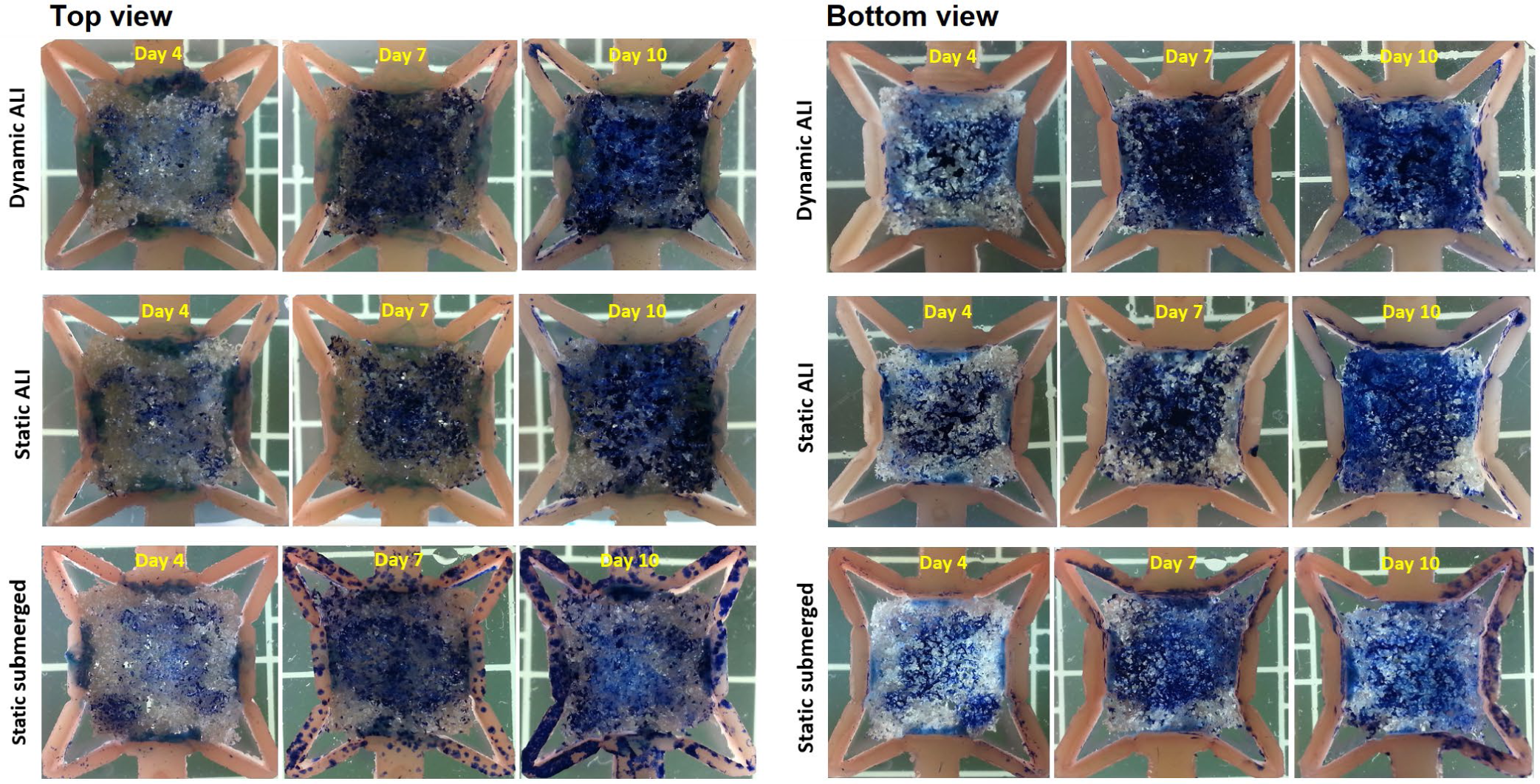
Higher magnification view of the stained constructs within the strain mechanism of the inserts (Left: top, Right: base). Note: the base of the constructs stained more intensely as passively seeded cells settled and established on the bottom side of the scaffolds

Cell colonies were observed on the strain mechanism and arms of the well insert in all groups from day 7 onwards. The greatest number of these ‘off-scaffold’ cells occurred in the static submerged group (Figure 4, Figure 5), suggesting that at lower oxygen tension further away from the surface of the culture medium (Figure 6), the cells displayed less preference for the porous scaffold compared to the actuation mechanism, which are composed of the same material. This hypothesis is further supported by how the off-scaffold cells in the submerged control group predominantly established at the top of the strain mechanism, i.e. the level closest to the surface of the culture media.

Higher retention of in-scaffold cells for both ALI groups suggests that oxygen tension is a key driver of phenotypic function, which is lost at a lower oxygen levels [37, 45]. This is aligned with *in vivo* conditions where lung cells function at atmospheric oxygen tension, which correlates to the surface of the culture medium i.e. the air-liquid interface of an *in vitro* culture environment. Furthermore, a smaller depth of fluid at the ALI compared to the bottom of the plate is a more physiological representation of the fluid conditions of the alveolar epithelium compared to submerged culture. Finally, displaced cells on the actuated groups preferentially established at the inner vertices of the strain mechanism (Day 10, Figure 5) i.e. the regions experiencing the most motion compared second to the scaffold; further indicating that mechanical stimulus has a positive effect on cellular behavior. Together, these results demonstrate that the preferred conditions for the respiratory cell line investigated are a combination of (i) atmospheric oxygen tension at an air-liquid interface, and (ii) mechanical stimulus, thus supporting our design hypothesis that air-liquid interface culture under physiological strain is more optimal for respiratory cell lines than conventional static submerged culture techniques.

#### D. Quantitative assessment of cell numbers in 3-D constructs

Total protein analysis was effective in providing a direct measure of cell numbers present within the 3-D scaffolds. A standard curve (R^2^ value > 0.995) was established from the absorbances of the BSA standards, from which the formula of the standard curve trend line and respective dilution factors were used to calculate total protein concentrations of the samples (Figure 7). All experimental groups had a similar cell number at day 4, and both the actuated and static air-liquid interface groups yielded higher cell numbers at days 7 and 10 compared to the static submerged culture condition (Figure 7). These trends directly corroborate the nuclei staining results for each time point, where both ALI group scaffolds were more darkly stained (i.e. contained higher cell numbers) than the static submerged group and had similar staining intensities (Figure 4, Figure 5). Lower total protein concentrations measured from the static submerged culture scaffolds directly correlates with lower numbers of on-scaffold cells revealed by nuclei staining, which indicated higher cell migration off the scaffold onto the PCL strain mechanism for the static submerged group compared to both ALI groups (Figure 4, Figure 5).

**Figure 7.**
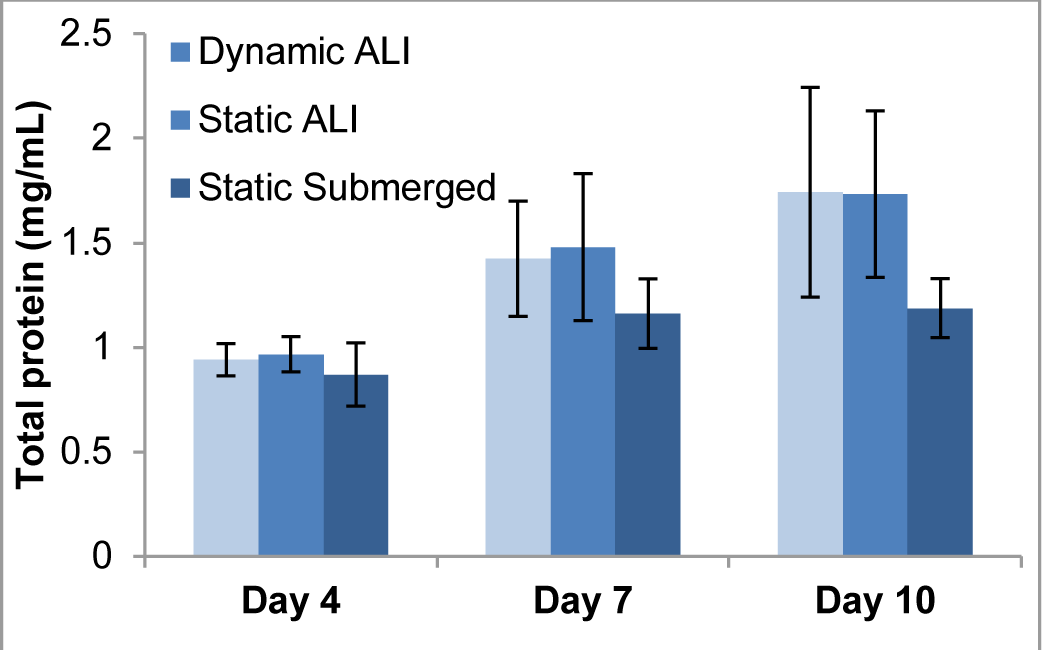
Protein concentrations of each experimental group expressed as mean ± SD, n = from 3 replicates

Interestingly, the static ALI group yielded a higher mean cell count than the actuated ALI scaffolds at day 4 (Figure 7). While not significant, further investigation is required to determine whether cells were shaken off the scaffolds under dynamic culture or followed different growth pathways as staining revealed different distribution patterns. Overall, the protein concentration assay provides a quantitative measure of cell proliferation and confirms the trends in cell growth observed with nuclei staining. We had previously investigated cell counting methods and encountered challenges including inconsistent and/or incomplete removal of cells, and cell damage from enzymatic digestion. In contrast, the total protein assay was found to be a straightforward and effective means of measuring cell numbers within thick scaffolds.

## V. Conclusion

In this study we present an *in vitro* system designed to provide physiological strain to 3-D cultures at an air-liquid interface. For proof of concept, a human lung tumor was modeled with NCI-H460 cells cultured on a 3-D synthetic scaffold the effect of applying mechanical stimulus was compared with static ALI culture and static submerged culture. High seeding efficacy was achieved by suspension within our scaffold well insert and distinctly different patterns in cellular growth were demonstrated between the conditions investigated. Significantly, scaffold cultures subjected to physiological strain and atmospheric oxygen tension exhibited the highest growth and most even cellular distribution, confirming the underlying biomimetic design rationale. Nuclei staining and total protein concentration analysis were readily adapted to meaningfully evaluate the 3-D constructs. Macroscopic imaging of the scaffold inserts enabled holistic assessment of cellular distribution throughout the culture environments and informed cellular behavior beyond in-scaffold growth. Total protein analysis allowed quantification of cell numbers, which corroborated trends observed from staining. While beyond the scope of this preliminary work, gene expression studies will further elucidate the differences in cellular behaviors observed. Overall, these promising results confirm our design rationale and underscore the importance of providing organ-specific conditions in engineering biofidelic tissues *in vitro*.

## Notes

### Competing Interest Statement

The authors have declared no competing interest.

